# Generation of a zebrafish neurofibromatosis model via inducible knockout of *nf2*

**DOI:** 10.1101/2024.04.23.590787

**Authors:** Ayyappa Raja Desingu Rajan, Yuanyun Huang, Jan Stundl, Katelyn Chu, Anushka Irodi, Zihan Yang, Brian E. Applegate, Marianne E. Bronner

**Author notes:** Equal contribution (Yuanyun Huang, Jan Stundl).

## Abstract

Neurofibromatosis Type 2 (NF-2) is a dominantly inherited genetic disorder that results from mutations in the tumor suppressor gene, neurofibromin 2 (NF2) gene. Here, we report the generation of a conditional zebrafish model of neurofibromatosis established by an inducible genetic knockout of *nf2a/b*, the zebrafish homolog of human NF2. Analysis of *nf2a* and *nf2b* expression reveals ubiquitous expression of *nf2b* in the early embryo, with overlapping expression in the neural crest and its derivatives and in the cranial mesenchyme. In contrast, *nf2a* displays lower expression levels. Induction of *nf2a/b* knockout at early stages increases the proliferation of larval Schwann cells and meningeal fibroblasts. Subsequently, in adult zebrafish, *nf2a/b* knockout triggers the development of a spectrum of tumors, including vestibular schwannomas, spinal schwannomas, meningiomas, and retinal hamartomas, mirroring the tumor manifestations observed in patients with NF-2. Collectively, these findings highlight the generation of a novel zebrafish model that mimics the complexities of the human NF-2 disorder. Consequently, this model holds significant potential for facilitating therapeutic screening and elucidating key driver genes implicated in NF-2 onset.

## Introduction

Neurofibromatosis Type 2 (NF-2) is an autosomal dominant disorder resulting from germline/mosaic mutations in the NF2 tumor suppressor gene, leading to multiple benign tumors in the nervous system and along peripheral nerves. Despite its benign nature, NF-2-associated tumors can lead to neurological deficits like early onset hearing loss, issues with balance, cataracts, seizures, pain, and problems with facial expressions. NF-2 tumors are primarily comprised of schwannomas, meningiomas, ependymomas, astrocytomas, and infrequently neurofibromas, retinal hamartomas, and intraorbital tumors (Campian and Gutmann, 2017; Coy et al., 2020; Ren et al., 2021; Tamura, 2021). While the estimated incidence of germline mutations in NF-2 is around 1 in 33,000, NF-2 also has one of the highest rates of mosaicism, with reports suggesting half of all individuals with NF-2 mutations have *de novo* genetic alterations (Chen et al., 2022). To date, the best treatment options for NF-2 are surgical removal, chemotherapy, and radiation therapy.

The neurofibromatosis (NF2) gene is a member of the ERM (ezrin, radixin, moesin) family of cell adhesion proteins and codes for the protein Merlin, which acts as a tumor suppressor. Merlin functions as a membrane-cytoskeleton linker that inhibits cellular proliferation via contact-dependent regulation of various signaling pathways, including WNT/β-catenin, Notch, Ras, Rac/Rho, TGF-β, Hippo, and receptor tyrosine kinases (Hamaratoglu et al., 2006; Okada et al., 2007; Zhang et al., 2010). During embryonic development, NF2 is highly expressed in various tissues. In the adult, its expression is predominantly observed in Schwann cells, meningeal cells, neurons, oligodendrocytes, mesothelium, optic neuroepithelial compartments, and lens fiber cells (Bakker et al., 1999; Moon et al., 2018; Toledo et al., 2018).

Modeling the plethora of phenotypes seen in NF-2 patients has been challenging, as biallelic knockouts of *Nf2* in mice are lethal due to failure to initiate gastrulation (McClatchey et al., 1997). Heterozygous/hemizygous knockouts of *Nf2* are cancer-prone and demonstrate a tumor spectrum that differs significantly from that observed in NF-2 patients; these do not develop schwannomas, a prominent feature of NF-2 (Giovannini et al., 2000; McClatchey et al., 1998). To avoid these issues, conditional knock-outs for *Nf2* have been generated in specific cell lineages. Conditional biallelic knockout of *Nf2* using P0-Cre transgenic mice leads to the development of schwannomas, cataracts, and tumors in tissues with neural crest-derived components. However, these mice do not develop vestibular schwannomas, the hallmark tumors of human NF-2 (Giovannini et al., 2000). Mice with conditional inactivation of the *Nf2* gene in leptomeningeal cells via subdural injection of *adCre* in *Nf2^flox2/flox2^*are prone to meningioma-genesis (Kalamarides et al., 2002). In contrast, conditional loss of *Nf2* using *Periostin-Cre* gives rise to vestibular schwannomas and Schwann cell hyperplasia in dorsal root ganglion (DRG) and proximal spinal nerve roots (Gehlhausen et al., 2015). However, each of these lines only forms tumors associated with that lineage and thus does not recapitulate the entire range of tumors in human NF-2 patients. Therefore, an animal model that better recapitulates the human disorder globally is still missing.

Here, we report the generation of an inducible *nf2* zebrafish knockout transgenic line using the CRISPR/Cas9 approach to model Neurofibromatosis type 2. Zebrafish embryos are transparent and develop *ex-utero*, easing the long-term visualization of both embryos and early larvae. Zebrafish offer several advantages, including rapid development, ease of intracranial imaging, and the ability to generate stable transgenic models that phenocopy human diseases, thus facilitating the study of lethal mutations. We show that *nf2* is expressed in the neural crest, meninges, and Schwann cells during early development. Loss of *nf2* results in aberrant proliferation of these cell types, eventually giving rise to schwannomas, meningiomas, cataracts, and abnormal pigmentation, thus creating a useful disease model in an organism accessible to imaging and genetic manipulation at low cost.

## Results

### Expression of *nf2a* and *nf2b* in the zebrafish embryo

Zebrafish have two orthologues of the human *NF2* gene, *nf2a* and *nf2b*. We first examined the expression of *nf2a* and *nf2b* transcripts in the 1-3 days post fertilization (dpf) zebrafish embryos using hybridization chain reaction (HCR), a highly sensitive *in situ* hybridization technique. Our results indicate that *nf2b* is the predominant paralog expressed in the early embryo at these time points. At 1 day post-fertilization (dpf), *nf2b* was broadly expressed in the cranial region, with notable signal in the forebrain, midbrain-hindbrain border, and the basal surface of the optic cup and lens. In addition, its expression overlapped with that of the neural crest marker, *sox10,* in cranial neural crest cells (Fig.1 A-C, A’-C’). The expression of *nf2a* was very weak compared to *nf2b,* with predominant expression in epithelial cells and some overlap with *sox10* expression (Fig.1 A, D-E, D’-E’). Next, we focussed on the expression of *nf2a* and *nf2b* in the cranial mesenchyme. The forkhead transcription factor *Foxc1* is a crucial regulator of cranial development and is expressed in the cranial mesenchyme before skeletogenic differentiation begins and is later restricted to the meningeal layers, cartilage primordium, and osteoblasts (Rice et al., 2003; Vivatbutsiri et al., 2008). Zebrafish have two orthologues of *Foxc1*, *foxc1a,* and *foxc1b*, while *foxc1a* is expressed highly in the cranial mesenchyme at 1dpf (Ferre-Fernández et al., 2022; Skarie and Link, 2009), *foxc1b* is expressed at later stages and labels the meningeal fibroblasts and the periocular mesenchyme. In transverse sections of the cranial region of 1 dpf embryos, we detected overlap between the transcription factor *foxc1a* and *nf2b* in the cranial mesenchyme (Fig.1 F-H, H’). In contrast to *nf2b, nf2a* was barely detectable at 1 dpf and displayed minimal overlap with *foxc1a* (Fig.1 I-K, K’).

**Fig 1:**
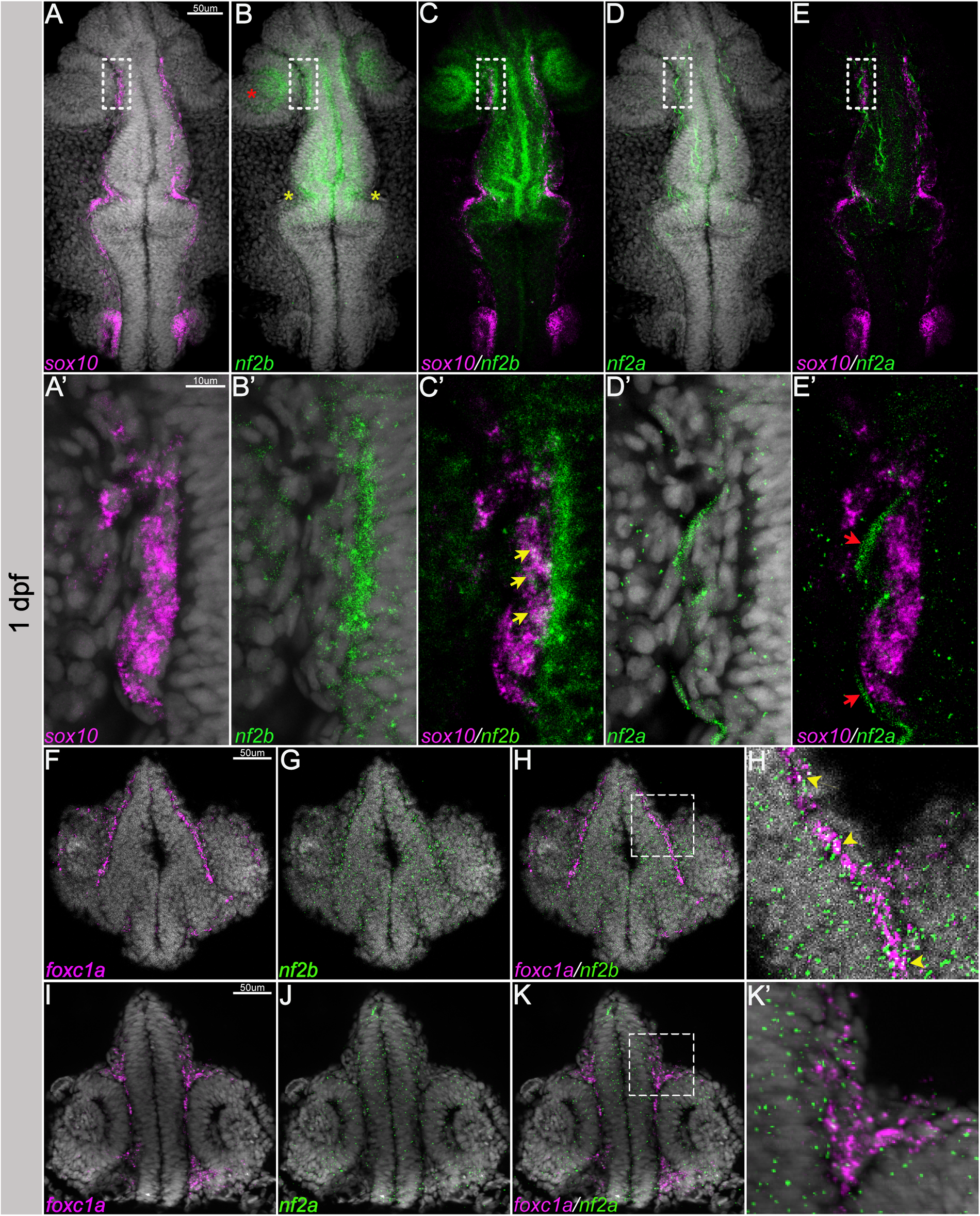
*nf2* is expressed in the cranial neural crest cells and mesenchyme. Multiplexed fluorescent mRNA in situ hybridizations by HCR reveals expression of (A-E) *nf2b*, *nf2a* and *sox10* in whole mount embryos at 24 hpf. (A’-E’) represents magnified image of the regions in white dotted box in (A-E) (yellow arrows represent overlap of *nf2b* and *sox10* expression, red arrow represents the ectodermal epithelial cell), (F-K) *nf2b*, *nf2a* and *foxc1a* in cryo-sectioned embryos at 24hpf. (H’, K’) represents zoomed of the regions in white dotted box in (H&K) (yellow arrowheads represent overlap of *nf2b* and *foxc1a* expression). Red asterisk - optic cup, yellow asterisk - midbrain-hindbrain border.

At 3 dpf, *sox10* expression is mainly observed in peripheral glia, spinal cord oligodendrocytes, and Schwann cells. At this stage, we noted several *nf2b* foci expressed in neural crest-derived Schwann cells along the posterior lateral line, labeled by *sox10* (Fig.2 B-D). We also observed a few *nf2a* foci in the Schwann cells (Fig.2 E-F). By 3 dpf, *foxc1b* is expressed in the meningeal fibroblasts and mesenchymal cells ventral to the forebrain region. We observed co-expression of both *nf2* paralogs and *foxc1b* in the meningeal fibroblasts, albeit at low levels for *nf2a* (Fig.2 G-L, I’ & L’).

**Fig 2:**
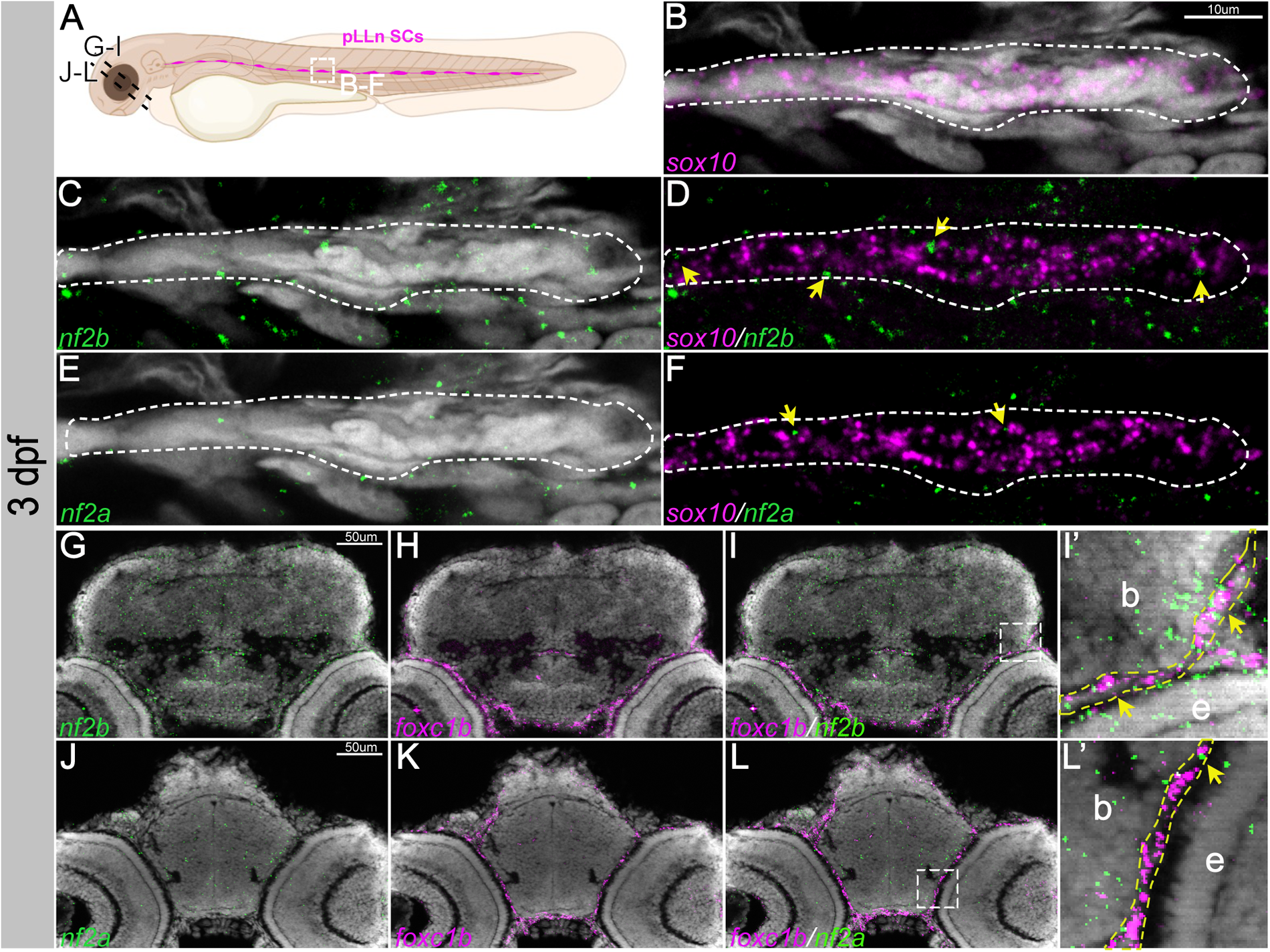
*nf2* is expressed in Schwann cells and meninges. (A) Schematic representing the posterior lateral line Schwann cells (pLLN SCs, dotted white box) and sectioned regions (dotted black lines) at 3dpf. Multiplexed fluorescent mRNA in situ hybridizations by HCR on whole mount embryos reveals expression of (B-F) *nf2b, nf2a* and *sox10* in a Schwann cell cluster at 3dpf. Yellow arrowheads indicate cells that have both *sox10* and *nf2* expression. Multiplexed fluorescent mRNA *in situ* hybridizations by HCR on cryo-sectioned embryos reveals expression of (G-L) *nf2b, nf2a* and *foxc1b* in the cranial meninges at 3dpf. (I’&L’) represents zoomed image of the regions in dotted white box in I and L respectively. Yellow dotted lines in (I’&L’) mark the meninges layer in close proximity to the brain and yellow arrowheads indicate cells that have both *foxc1b* and *nf2* expression. b-brain, e-eye.

Our results indicate that the zebrafish *nf2* paralogs are expressed broadly during early development, notably in the cranial neural crest, cranial mesenchyme, optic cup, and lens. This is consistent with observations of *Nf2* promoter and mRNA expression in mouse embryos (Akhmametyeva et al., 2006; Huynh et al., 1996). Additionally, we detected the expression of *nf2* paralogs in the developing Schwann cells and meningeal fibroblasts. Interestingly, *nf2b* is much more highly expressed in most tissues than *nf2a*, indicating that it is the dominant paralog. However, *nf2a* but not *nf2b* is expressed in epidermal cells, suggesting that these paralogs may also have some cell type-specific function.

### Generation of an inducible *nf2* knock-out line

To establish a neurofibromatosis type 2 model, we used CRISPR-Cas9 to target zebrafish *nf2a and nf2b* (collectively referred to as *nf2*). First, we tested guide RNAs (gRNAs) targeting both the paralogs of zebrafish *nf2*. After determining the knockout efficiency, four gRNAs (two for each paralog) were selected for cloning into pU6x:gRNA vectors for constitutive gRNA expression. The *nf2* gRNAs containing pU6x:gRNA vectors were cloned into pGGTol2-LC-Dest-4sgRNA vector and injected into wild type 1-2 cell zebrafish embryos with *Tol2* transposase mRNA for genomic integration. Embryos were sorted at 3dpf for cerulean expression in the lens and grown to adulthood. Once stable F1/F2 lines were established, we used semi-quantitative RT-PCR to examine gRNA expression (Suppl Fig. 1A). As expected, we observed expression of the gRNAs only in the lens-cerulean-positive embryos compared to cerulean-negative embryos.

Next, to test the mutagenesis efficiency of the *Tg(pU6x:nf2-4sgRNA)* expressing stable lines, we crossed the fish with the *Tg(hsp70:loxP-mCherry-STOP-loxP-cas9)* line, referred to as the *Tg(HOTCre:cas9)* line. This allows the induction of Cas9 expression by heat shock in a Cre-dependent manner in the presence of the gRNAs (Yin et al., 2015) (Fig. 3A illustrates the knock-out strategy). Using a semi-quantitative T7 endonuclease assay, we observed all four gRNAs could elicit mutations *in vivo* at almost 80-95% efficiency (Suppl Fig. 1 B). We performed PCR sequencing for the *nf2a* and *nf2b* target regions and observed CRISPR-Cas9-induced modifications in 70-95% of the reads (Fig. 3 B-C). Using Western blot analysis, we observed the concomitant loss of the Nf2 protein in the CRISPR-Cas9-induced animals (Suppl Fig. 1C).

**Fig 3:**
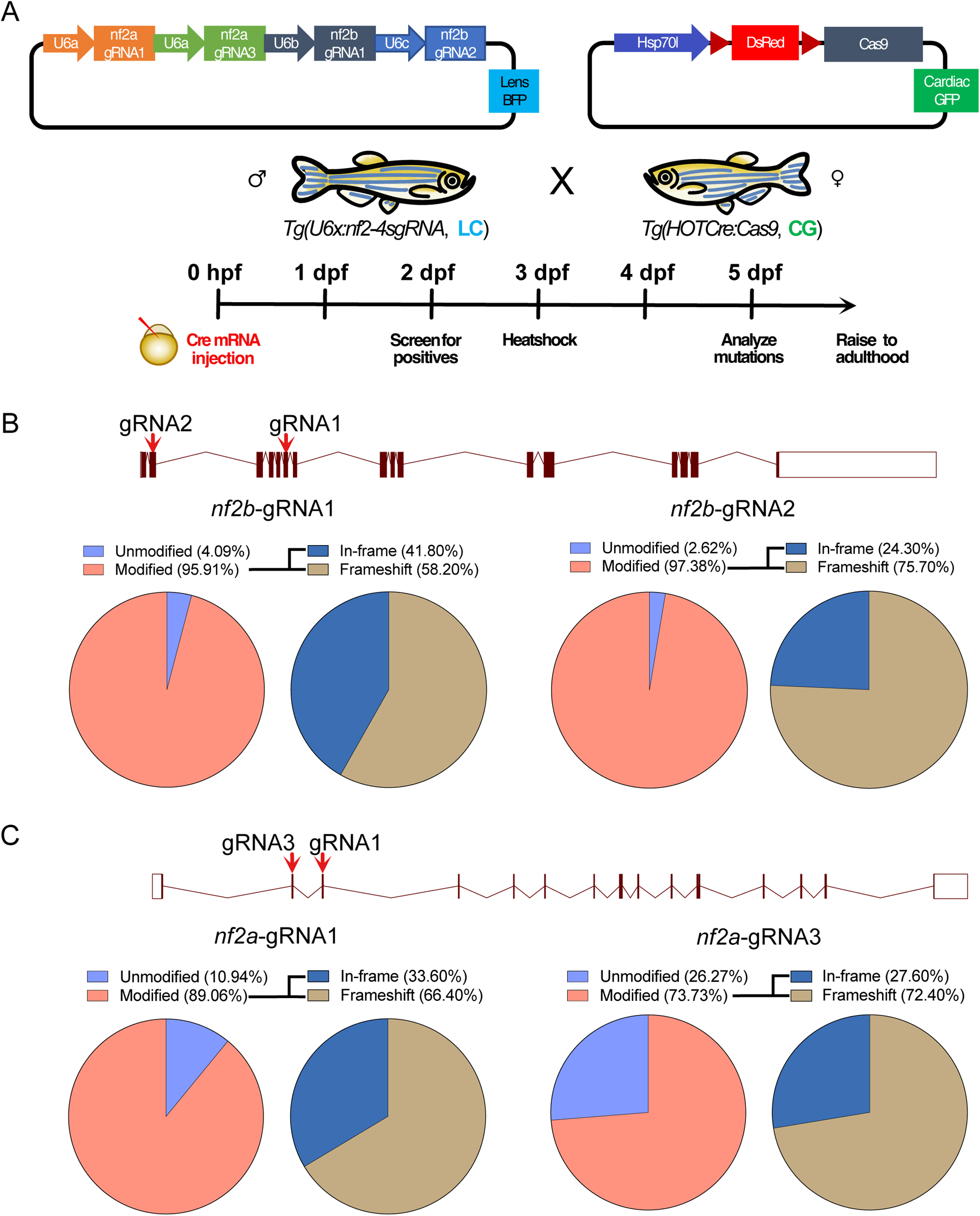
Strategy for generation of conditional knockouts for *nf2b* and *nf2a.* (A) Schematic of strategy used to generate *nf2* knockouts. Schematic of target regions and Pie charts represent percentage of mutated reads in (B) *nf2b*, (C) *nf2a* knockout embryos. LC - lens cerulean, CG - cardiac green fluorescent protein.

Biallelic knockouts of *Nf2* mice die early during embryonic development due to a failure to initiate gastrulation because of an absence of organized extraembryonic ectoderm (McClatchey et al., 1997). In contrast, tissue-specific *Nf2* knockouts give rise to tumors only in the associated lineages. Our strategy circumnavigates these issues by enabling tight control of the timing of Cas9 expression and the resulting mutations via heat shock to avoid disrupting *nf2* at early developmental stages. We tested the survival of mosaic *nf2* KO embryos upon induction of Cas9 expression on days 1, 3, and 5 of development. Day 3 induction of Cas9 resulted in the maximum number of survivors until day 15 of development compared to day 1 and day 5 induction. Nevertheless, induction of *nf2* mutation at early embryonic and larval stages largely led to death, with ∼20-25% survivors for day 3 induction (Suppl Fig. 1D).

### Effects of *nf2* knockout of neural crest derivatives, like Schwann cells, melanocytes, and meninges

Given that zebrafish *nf2* is expressed in neural crest cells, cranial mesenchyme, schwann cells, and the meninges, we tested the effect of *nf2* knockout on proliferative ability in these cell types. First, we compared meningeal cell proliferation at 3dpf between Cas9-only controls and *nf2* conditional knock-outs (heat shock at 1 dpf). We used *igfbp2a* as a marker to label the meninges and found increased proliferation in the meningial cells after nf2 knock-out (Fig. 4 B-G, E’-G’). Quantitation of this effect demonstrates a significant increase in meningeal pH3 staining after the loss of *nf2* (Fig. 4 H) Next, we utilized the transgenic line *Tg(*-7.2 *sox10:mRFP)* to visualize Schwann cells along the posterior lateral line at 3dpf and 5dpf. While we did not see any difference in the schwann cell number at 3dpf between *nf2* knockout and Cas9-only controls (Fig. 5 B, C, F & G)), we observed a striking increase in the numbers and thickness of Schwann cell clusters along the posterior lateral line in *nf2* knockout larvae (Fig. 5 H-J) compared to the Cas9-only controls (Fig. 5 D, E) at 5dpf, an observation strikingly similar to zebrafish *nf1* knockouts (Shin et al., 2012).

**Fig 4:**
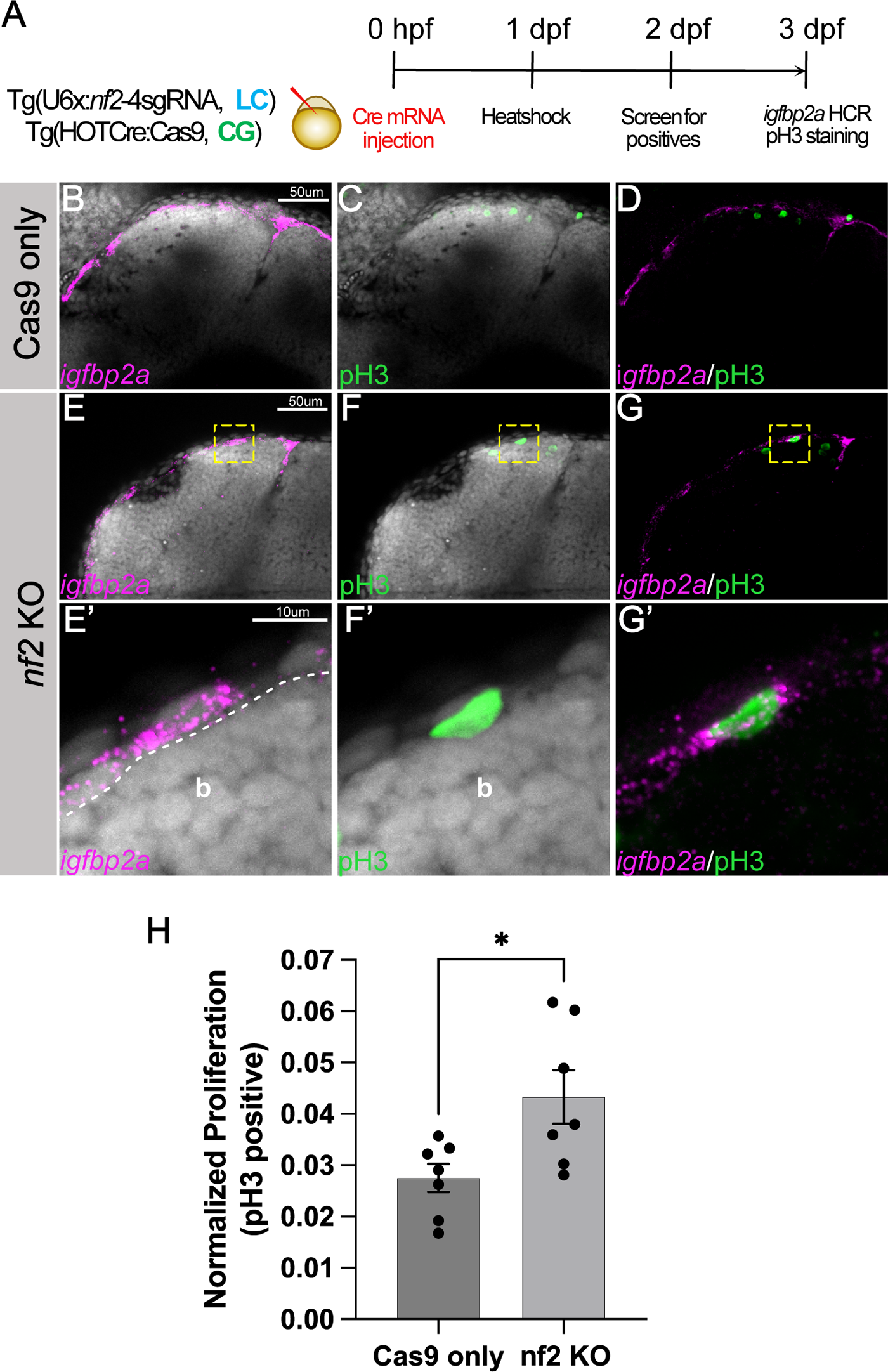
*nf2* knockout leads to increased meningeal proliferation. (A) Schematic representing the experimental strategy. Confocal images of the cranial region of 3 dpf Cas9 only (B-D) and *nf2* KO (E-G) embryos show the overlap of meningeal marker *igfbp2a* and proliferation marker phospho-histone3 (pH3). (E’-G’) Yellow dotted boxes shows the magnified region of the overlap in the *nf2* KO embryos. (H) Bar plot showing quantification of normalized proliferation (p-values: *, p=0.0197, each dot represents data of one larva). Each dataset was compared with Student’s t-test using GraphPad Prism. White dotted line marks the boundary between brain and meninges.b - brain.

**Fig 5:**
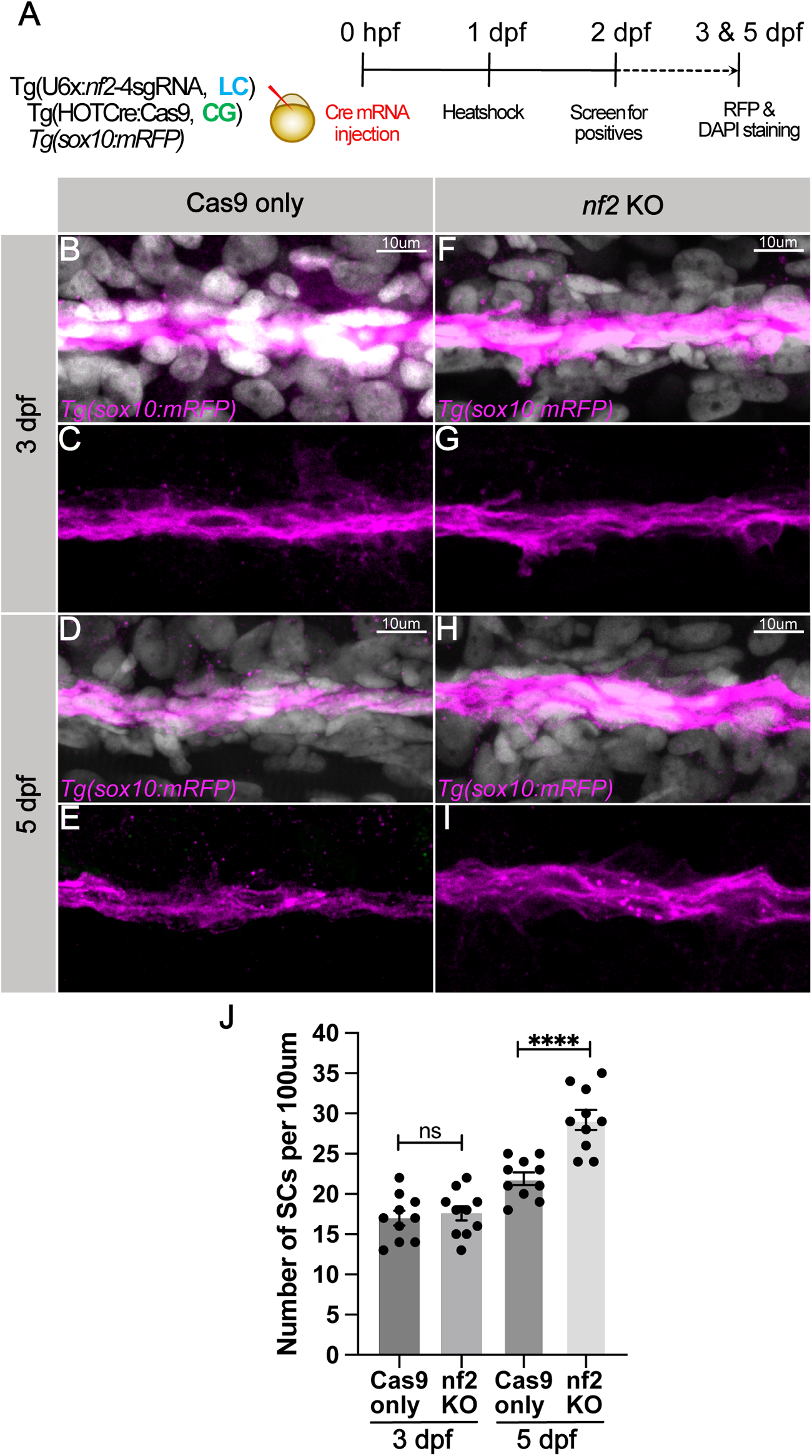
*nf2* knockout leads to Schwann cell hyperplasia. (A) Schematic representing the experimental strategy. Confocal images of the posterior lateral line schwann cells (pLLn SCs) of Cas9 only controls (B-E) and *nf2* KO (F-I) embryos at 3dpf and 5dpf. pLLn SCs are labelled by *Tg(sox10: mRFP)*[magenta] and counterstained by DAPI [grey]. (J) Bar plot showing quantification for the Sox10-positive SCs along the pLLN (p-values: ns=not significant; ** p<0.0001, each dot represents data of one larva). Each dataset was compared with Student’s t-test using GraphPad Prism.

NF-2 patients may frequently present with cafe-au-lait macules, which are hyperpigmented regions in the skin. Consistent with this, we observed striking hyperpigmentation in *nf2* knockouts compared with Cas9-only controls at 3dpf (Fig. 6 B, C, F, G). At this stage, most of the Cas9-only control melanocytes are settled in the dorsal/midline/ventral pigmentation pattern; however, we noted that melanocytes in *nf2* knockouts melanocytes are not completely patterned and appear to still be migratory (Fig. 6 F; red arrows). In addition, we observed a marked increase in melanocyte density in the cranial regions (Fig. 6 F; green arrows). At 6dpf, we observed dramatic hyperpigmentation of the head, reduced body length, small eyes, and impaired inflation of the swim bladder in *nf2* knockouts (Fig. 6 H, I). Taken together, the results suggest that the loss of *nf2* in zebrafish affects many of the same cell types that are prone to tumorigenesis in human NF-2 patients.

**Fig 6:**
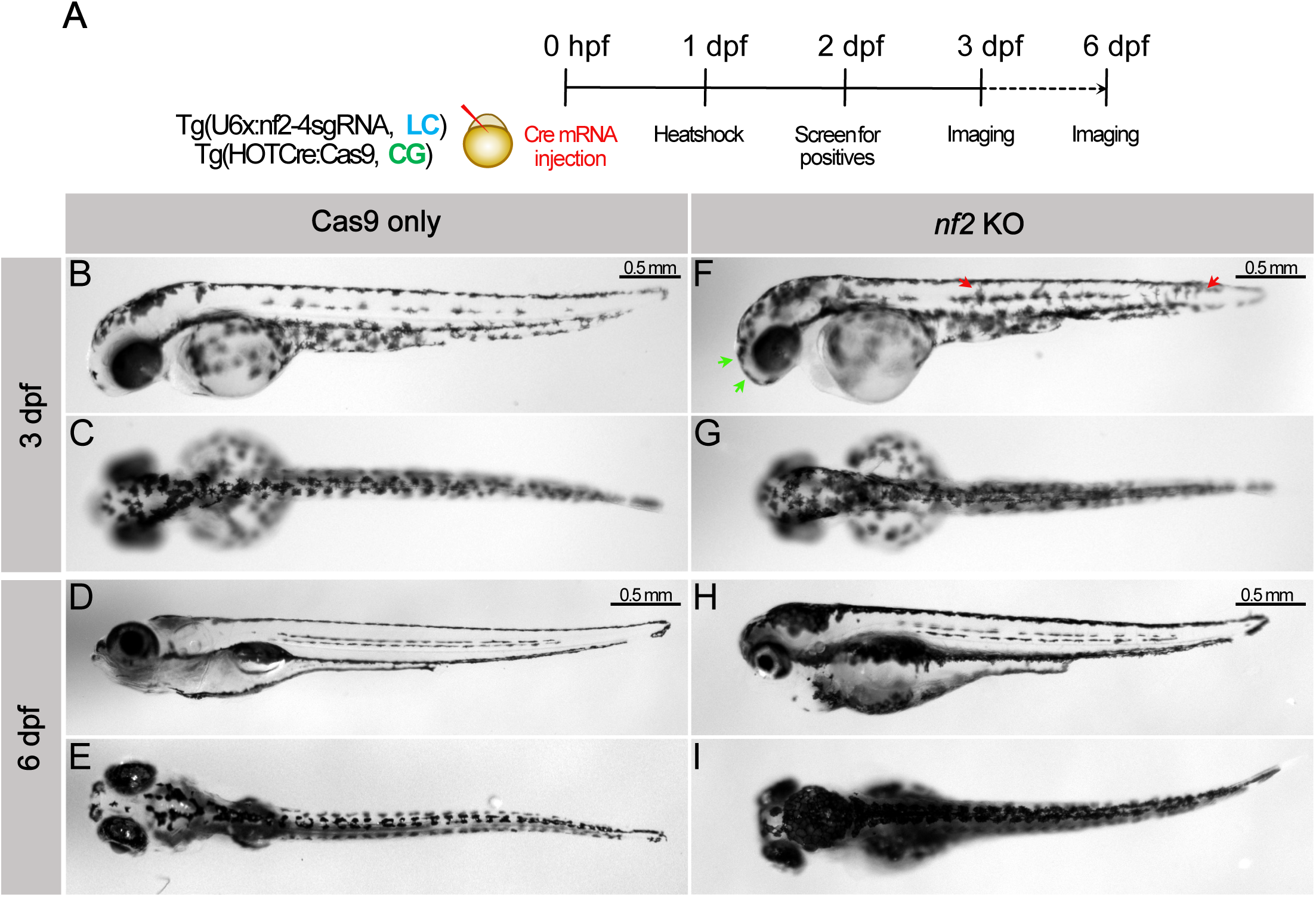
Knockout of nf2 leads to hyperpigmentation. (A) Schematic representing the experimental strategy. Lateral and dorsal bright-field images of Cas9 only control (B-E) and *nf2* knockout (F-I) embryos at 3dpf and 6dpf. Green arrows - increased melanophores in cranial region, red arrows - migratory melanophores.

### Tumor formation in adult NF2 knockout zebrafish

Given that most embryos die after heat-shock-mediated Cas9 induction during the first few days of development, for long-term survival that would allow analysis of tumor formation, induction was conducted in 3-6-month-old fish. Zebrafish were allowed to develop 1-10 months after the induction. At these time points, we noted issues with balance, cataracts, hyperpigmented regions, and conspicuous tumors. Interestingly, loss of balance and issues with swimming were some of the first phenotypic manifestations observed in the *nf2* knockouts. Zebrafish *nf2* knockouts had tumors of several subtypes, primarily comprised of vestibular and spinal schwannomas, meningiomas, retinal hamartomas, and ependymomas.

To characterize tumor morphology, adult fish were fixed and sectioned in the transverse plane. Sections were stained with hematoxylin/eosin and examined at the brain or spinal cord level and surrounding tissues. In Cas9-only control animals, we did not observe any aberrant growth of the involved tissues (Fig. 7 B, D, F, H, J & L ). In the *nf2* knock-outs, we observed transitional meningiomas (containing fibroblastic and meningothelial components). In addition, some of these meningiomas penetrated the cranium and appeared metastatic (Fig.7 C, C’). When focusing on the cranium, the Cas9-only control fish appeared normal (Fig. 7 D), whereas the *nf2* knockouts displayed thickening of the bones (Fig. 7 E, E’) (Li et al., 2018).

**Fig 7:**
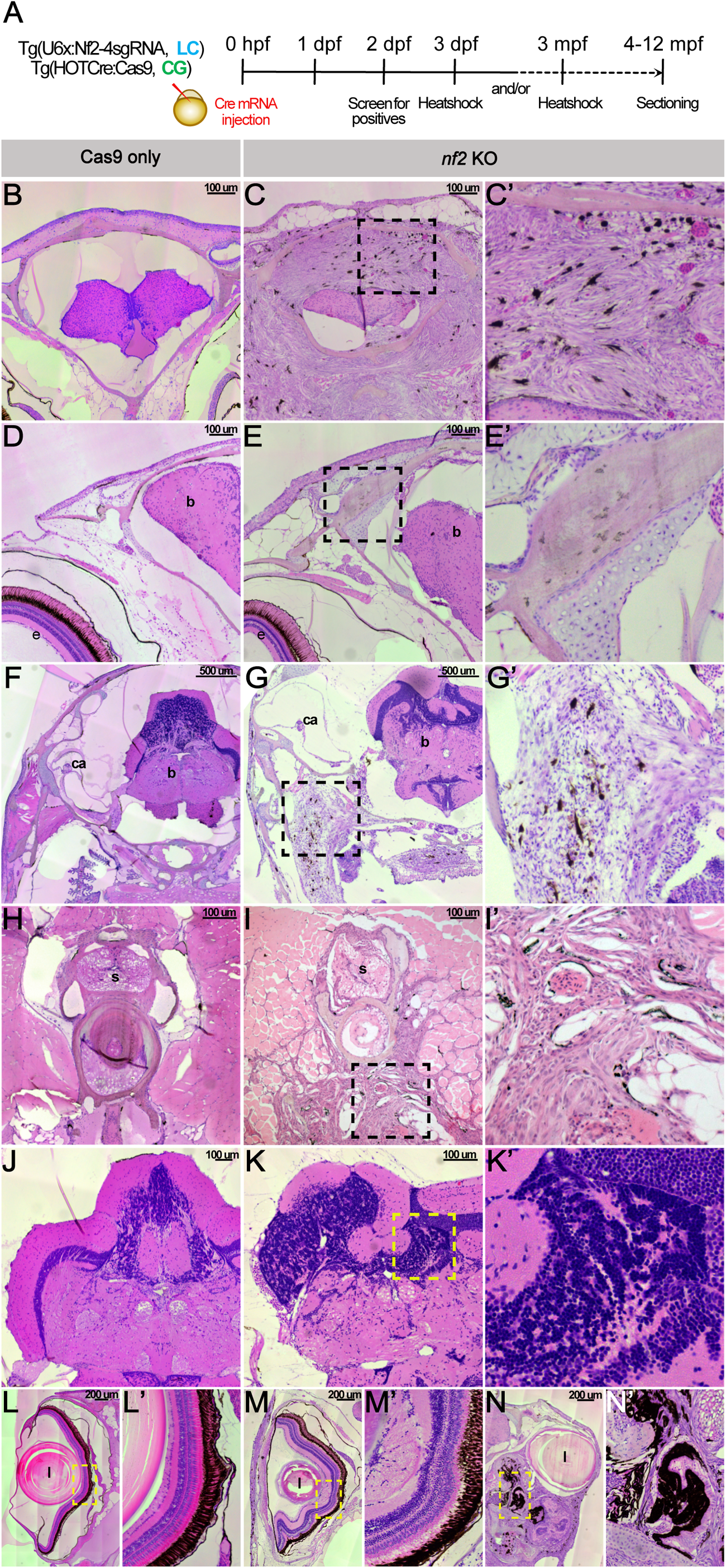
*nf2* knockout leads to tumors in adult zebrafish. (A) Schematic representing the experimental strategy. Transverse sections of *nf2* KO zebrafish adult displays meningioma (C-C’), skull bone thickening (E-E’), vestibular Schwannoma (G-G’), trunk Schwannoma (I-I’), ependymoma (K-K’), epiretinal membrane (M-M’) and retinal hamartoma (N-N’). Cas9 only sections (B,D,F,H,J & L) for comparison. Black and yellow dotted boxes represent the zoomed regions. Majority of the tumor samples display abundant melanocytes within the tumors. e-eye, b-brain, i-semicircular canal of inner ear, l-lens and s-spinal cord.

Bilateral vestibular schwannomas are one of the primary diagnostic features of neurofibromatosis type 2. While the auditory nerves of controls appeared normal (Fig. 7 F), by contrast, vestibular schwannomas were observed in the *nf2* knockout animals (Fig. 7 G, G’). At the spinal cord level, we noted schwannoma-like tumors that appeared to have aggressively metastasized into muscle tissues in the *nf2* knockouts (Fig. 7 I, I’). We also observed cranial ependymomas (Fig. 7 K, K’). Finally, we looked at the sections of the eye in the Cas9-only controls, which are arranged into distinct retinal layers similar to the human retina (Fig. 7 L-L’) (Richardson et al., 2017). The *nf2* knockouts displayed a distorted morphology with epiretinal membranes and loss of distinct layers (Fig. 7 M-M’). Additionally, we observed some knockouts with severe retinal hamartomas (Fig. 7 N-N’).

Ideally, we would like to recognize tumors in a noninvasive manner. To this end, we subjected 5 to 10-month-old Cas9-only control and *nf2* knockout animals to an analysis by optical coherence tomography (Suppl Fig. 2). The results demonstrate increased telencephalic ventricular volume in the *nf2* knockouts compared with Cas9-only controls. A possible explanation is that the tumors may obstruct cerebrospinal fluid flow, resulting in enlarged intraventricular space that can be detected in the intact animal. This is consistent with observations of hydrocephalus resulting from cerebrospinal fluid blockage in some NF-2 patients (Tanrıkulu and Özek, 2019).

## Discussion

Neurofibromatosis type II (NF-2) is an inherited condition that increases the risk of developing specific nervous system tumors such as bilateral vestibular schwannomas, multiple spinal and peripheral schwannomas, meningiomas, and ependymomas. NF-2 is caused by inactivating mutations in the *NF2* gene, which may be germline or somatic. Two clinical forms of NF-2 have been historically documented. The Wishart phenotype represents a more aggressive manifestation of the condition, characterized by the development of multiple neoplasms in patients under 20 years old, with rapid progression of lesions. On the other hand, some patients may display a less severe phenotype known as the Gardner phenotype, characterized by fewer slow-growing tumors that typically appear later in life. It is now understood that the specific type of alteration in the NF2 gene primarily influences the severity of the disease spectrum. Patients with truncating alterations that deactivate NF2 tend to experience a more severe disease course, while those with missense loss-of-function mutations generally have a milder disease progression (Halliday et al., 2017).

To date, existing mouse models for NF-2 have failed to fully recapitulate the human phenotype. While homozygous loss of *Nf2* causes early mortality (McClatchey et al., 1997), hemizygous or heterozygous loss of *Nf2* results in a tumor spectrum different than that seen in the human counterparts (McClatchey et al., 1998). Tissue-specific conditional knockouts produce tumors in select tissues rather than the whole range of tumors found in patients (Gehlhausen et al., 2015; Giovannini et al., 2000). In recent years, zebrafish has become a robust model organism for cancer research due to their rapid embryonic development, high fecundity, amenability to genetic manipulation, drug treatments, and transparency throughout early developmental stages, allowing for powerful in vivo imaging (Patton et al., 2021; Roy et al., 2024; Shin et al., 2012; White et al., 2013). Moreover, the results obtained from zebrafish models can be translated back to humans due to the highly conserved nature of these cancer-related programs. Thus, the zebrafish serves as a non-mammalian vertebrate organism that represents a cost-effective model to study the effects on tumor formation of loss of genes like *NF2*.

Our study found a broad range of phenotypes in 5-10-month-old zebrafish after mutagenesis of *nf2* in 3-month-old fish—these range from meningiomas, vestibular schwannomas, cataracts, retinal hamartomas, and ependymomas. While many animal models require the knock-out of a gene in a sensitized background (Shin et al., 2012), we find that the *nf2* mutations alone appear sufficient in wild-type fish to produce tumors. A likely explanation for these observations is the inducible nature of *nf2* mutagenesis, wherein we can elicit biallelic mutations (a prerequisite for NF-2-related tumorigenesis) at the desired time of the animal’s lifespan. We demonstrate that zebrafish *nf2* is expressed in cell types, such as cranial neural crest, Schwann cells, and meningeal fibroblasts, during early development that later are associated with NF-2 tumors. Additionally, the knockout of *nf2* during early development results in the hyperproliferation of these cell types.

While the tumors arising in NF-2 seem to be in different anatomical locations, one likely cell of common origin for NF-2 tumors is Schwann cell precursors (SCPs). These are neural crest-derived cells that persist along peripheral nerves through adulthood and can migrate, differentiate, and dedifferentiate under appropriate conditions. While able to form Schwann cells, SCPs are often multipotent and can differentiate into pigment cells and a wide range of neural crest-derived cell types (Solovieva and Bronner, 2021). Their broad developmental potential correlates with the broad range of tumor phenotypes in NF-2 patients, including spinal, peripheral, and cranial nerve tumors (Gehlhausen et al., 2015).

A puzzling observation from our study and earlier work is that even though *NF2* is widely expressed during embryonic development, why does its inactivation predispose tumors in specific tissues? According to Knudson’s two-hit hypothesis, it is believed that NF-2-associated tumors occur due to additional somatic genetic alterations in susceptible cell populations, leading to the bi-allelic loss of function of NF2 (Woods et al., 2003). Another likely explanation could be the mechanosensitivity of the involved tissue types. The tumors prevalent in NF-2, like meningioma, schwannoma, retinal hamartoma, et.c, arise from precursor cell types like meningeal fibroblasts, schwann cells, and retinal cells, respectively. These cells are known to be present as stratified layers (e.g., meninges and retina) or tightly woven in peripheral nerves (e.g., Schwann cells). Such arrangement can lead to changes in the nuclear to cytoplasmic connections and facilitate the aberrant transport of nuclear effector/oncogenes in the *NF2* mutant background. One such example is the protein YAP, which is known to aid *NF2-*mediated tumorigenesis (Guerrant et al., 2016; Laraba et al., 2022; Oh et al., 2015; Szulzewsky et al., 2022). While several studies have explored the functional role of YAP in combination with NF2 in tumorigenesis, the mechanobiological aspect of their interaction is yet to be decoded. Recently, Alasaadi *et al*., 2024 demonstrated the role of YAP in neural crest competency via hydrostatic pressure. Thus, it is possible that tissue mechanics may interplay with signaling pathways to regulate tumor induction in NF-2.

In summary, we describe a new zebrafish model for conditional inactivation of the *nf2* gene. Since our inducible model can undergo conditional inactivation of *nf2* over various ages, this model can potentially recapitulate neurofibromatosis onset at different stages of life. Moreover, it holds the promise of testing the effects of therapeutic agents. Alternatively, this model will enable testing the effect of large-scale chemical libraries on the ability to ameliorate the phenotypes of NF-2. The results demonstrate the utility of this model for recapitulating a broad range of phenotypes associated with NF-2 and closely mimic human disease. The accessibility, ease of manipulation, availability of genetics, and facility of imaging promise to make this an extremely useful model for further exploration of tumor ontogeny and assaying means of treatment.

## Material and Methods

### Zebrafish lines

Zebrafish (*Danio rerio*) were maintained at 28°C, with adults on a 14-hour light/10-hour dark cycle. All zebrafish work complied with the California Institute of Technology Institutional Animal Care and Use Committee (IACUC). Transgenic lines used in this study were the ABWT (ZIRC), *Tg(HOTCre:Cas9)* (a Kind gift from Dr Wenbiao Chen), *Tg(-7.2kb-sox10:mRFP),* and the *Tg(U6x:nf2-4sgRNA)* line (generated at the Bronner laboratory, Caltech).

### Hybridization chain reaction and Immunohistochemistry

HCR v3.0 was performed according to the protocol suggested by Molecular Technologies zebrafish with several modifications. Briefly, methanol-fixed embryos were rehydrated by a series of methanol/PBS-Tween solutions (2 x 100%, 75%, 50%, 25%; every step 15 min), washed in PBS Tween (2 x 10 min), depigmented by a bleaching mix-solution (Formamide, 20x SSC, 30% hydrogen peroxide, and distilled water) under the direct light, washed in PBS-Tween (10 min), treated with proteinase-K (20ug/ml) according to the age of the larvae, washed in PBS-Tween (2 x 10 min), post-fixed in 4% PFA (20 min), washed in PBS-Tween (3 x 10 min), pre-hybridized in 30% probe hybridization buffer at 37°C (60 min), and incubated with probes (2 - 4 μl of 2 μM stock per probe mixture) in probe hybridization buffer at 37°C overnight. All probes, hairpins, and buffers were designed and ordered through Molecular technologies (https://www.molecularinstruments.com). The samples for histological analysis were embedded in agarose/OCT, sectioned (12um), and counterstained with DAPI. All photographs were taken with Zeiss AxioImager.M2 equipped with an Apotome.2.

Phospho-histone 3 (pH3) immunostaining was performed on either HCR-processed or methanol-fixed embryos for proliferation assays. Briefly, HCR-processed embryos were transferred to 1X PBS-Triton X-100 (3 x 10 min), or methanol-fixed embryos were rehydrated by a series of methanol/PBS-Triton X-100 solutions (2 x 100%, 75%, 50%, 25%; every step 15 min), blocked with 10% donkey serum - 1X PBS-Triton X-100 for 4 hours at room temperature. The embryos were incubated in anti-phospho-histone 3 antibody overnight at 4^0^C. The next day, the samples were washed in PBS-Triton-X100 (4 x 30 min) at room temperature and incubated in a secondary antibody (A21202, ThermoFisher Scientific) for 4 hours at room temperature. The samples were then rewashed in PBS-Triton-X100 (4 x 30 min) at room temperature. All whole-mount images were imaged with a Zeiss LSM 900 confocal microscope.

### CRISPR-Cas9 strategy

Guide RNAs for zebrafish *nf2a* and *nf2b* (Suppl Table 1) were designed using CHOP-CHOP (https://chopchop.cbu.uib.no/). Guide RNAs targeting zebrafish *nf2* were validated and cloned in the pU6(x):sgRNA#(x) vectors. All four guides were cloned into the final destination vector pGGDestTol2LC-4sgRNA (*nf2*-4sgRNA). The pU6(x):*nf2*-4sgRNA vector and Tol2 transposase mRNA were injected into ABWT 1-2 cell embryos. Embryos were sorted at 3dpf for cerulean expression in the lens and grown to adulthood. Two independent stable lines were established for the pU6(x):*nf2*-4sgRNA transgene (Suppl Fig. 1 A-B). For generating *nf2* knockouts, *Tg(U6x:nf2-4sgRNA)* fish were crossed with *Tg(hsp70:loxP-mCherry-STOP-loxP-cas9)* referred to as *Tg(HOTCre:Cas9)* fish. 10-20 pg of Cre recombinase mRNA was injected into the embryo. Heat shock was performed by adding 40^0^C E3-water, and incubation at 38^0^C for 15 minutes and 30 minutes was applied to embryos or adults, respectively, at the indicated stages.

### T7 Endonuclease assay and Premium PCR sequencing

Genomic DNA was isolated from control and *nf2* knockout embryos following the protocol described in https://zfin.org/zf_info/zfbook/chapt9/9.3.html. Guide RNA target regions for *nf2a* and *nf2b* were amplified using the primers described in supplementary table 1. T7 endonuclease reaction was set up according to (Lingeman, 2017). The digested products were then electrophoresed in a 2% agarose gel. Purified PCR products were sent for premium PCR sequencing to Primordium Labs (https://www.primordiumlabs.com/). At least 10,000 high-quality reads were analyzed for each sample using CRISPResso2 (Clement et al., 2019).

### RNA isolation, cDNA synthesis, and PCR

RNA was isolated from zebrafish embryos using Nucleospin Triprep (Macherey Nagel, 740966) according to the manufacturer’s protocols. cDNA was synthesized using the Superscript III First-Strand cDNA Synthesis Kit (ThermoFisher Scientific, 1800051). PCR was carried out using the primers described in Supplementary Table 1. PCR products were electrophoresed in a 1.5% agarose gel.

### Protein extraction and western blotting

Zebrafish embryos were dechorionated and deyolked in an ice-cold ringer solution. Approximately 100 embryo bodies were resuspended in 50 ul of NP40 lysis buffer (ThermoFisherScientific; FNN0021). The NP40 lysis buffer was supplemented with a 1X protease inhibitor cocktail (SIGMA; 05892970001). The embryo bodies were crushed using a pestle homogenizer and kept on ice for 15 minutes. The embryo bodies were homogenized again and kept on ice for 15 more minutes. The sample was centrifuged at 13,800 rcf for 20 minutes at 4^0^C. The soluble fraction of cell lysate was collected in a fresh tube. The protein concentration in the soluble fraction was quantified using bicinchoninic acid (BCA) protein estimation kit using known concentrations of bovine serum albumin (BSA) as standard. 30-50 μg of the protein was run on 10 % SDS PAGE gels. Proteins were separated by using 25 mA per gel in the electrophoresis buffer. The resolved samples were transferred to 0.2um Immobilon-FL PVDF membranes (Millipore) (activated in absolute methanol for 30 seconds) using wet transfer (in transfer buffer containing 14.4 g Glycine, 3.03 g Tris, and 20% methanol). The transfer was carried out for 90 minutes at 300 mA at 4°C, and membranes were blocked with 5% skim milk in 0.1% TBST for 1 hour at RT. Subsequently, the blot was incubated with 1:1000 dilution of anti-Merlin antibody overnight at 4°C. Subsequent to incubation with the primary antibody, the membrane was washed with 0.1% TBST and incubated with corresponding secondary antibody conjugated to Horse Radish Peroxidase (HRP) at 1:10000 (GE life sciences; NA934) dilutions for one hour at room temperature. The membrane was washed with 0.1% TBST and developed using ECL reagent (Millipore; WBKLS0500). Immunoblots were imaged using conventional chemiluminescent immunoblotting

### OCT imaging

OCT imaging of the zebrafish brain was conducted using a custom OCT system, with details of the system described in a previous paper (Kim et al., 2019). Briefly, the OCT system comprises a swept source operating at 1310 nm, offering a bandwidth of ∼ 93 nm and a sweep rate of 100 kHz. This setup affords an axial resolution of ∼ 18 μm (in air) and a lateral resolution of ∼ 30 μm. The euthanized zebrafish were placed in a petri dish with a foam holder to secure their position. The custom OCT system, integrated with a camera and an indicator laser, assisted in determining the imaging position of the zebrafish. Three-dimensional OCT images were acquired, covering an imaging area of 9 x 9 mm.

### Zebrafish adult head histology

The adult zebrafish heads for histological analysis were prepared as described in (Stundl et al., 2023); briefly, the samples were rinsed in distilled water, decalcified in Morse’s solution, and embedded into the JB4 resin (prepared according to the manufacturer’s instructions, Sigma-Aldrich) at room temperature overnight. The next day, the infiltration solution was replaced by an embedding solution (prepared according to the manufacturer’s instructions), placed into an embedding mold (Polyscience), and transferred to the vacuum chamber, which accelerated the polymerization (∼3 hours). The resin block was sectioned at 7 μm, and sections were stained with Mayer’s hematoxylin.

## Acknowledgements

We thank the Beckman Institute Biological Imaging Facility of Caltech for technical assistance with microscopy experiments. We thank the Wenbiao Chen lab for generously supplying transgenic fish *Tg(HOTCre:Cas9)*. We thank Justin Yip, Ryan Fraser, and David Mayorga for their help with fish facility maintenance. We thank Johanna Tan-Cabugao and Constanza Gonzales for their technical assistance. We would also like to thank all our fishes for providing embryonic and adult material for our research.

## Author contributions

Conceptualization—D.A.R. and M.E.B. Methodology—D.A.R. and M.E.B. Investigation—D.A.R, Y.H, J.S, K.C, and A.I; Z.Y, D.A.R, and B.E.A. performed and analyzed OCT data. Data curation—D.A.R. Visualization—D.A.R and M.E.B. Writing—original draft preparation—D.A.R. and M.E.B. Supervision—M.E.B. Funding acquisition—M.E.B.

---

**Supplementary Table 1.**
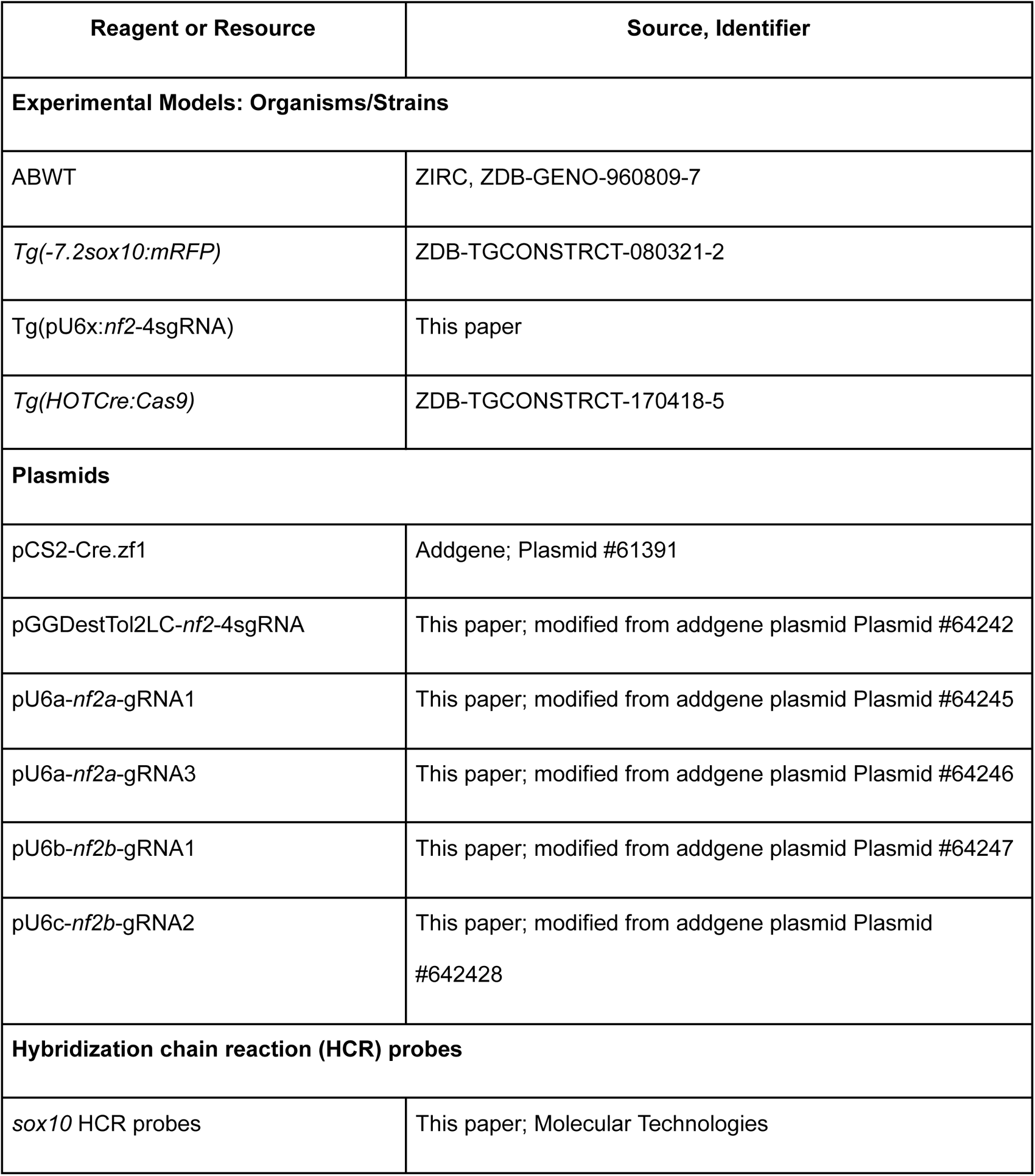
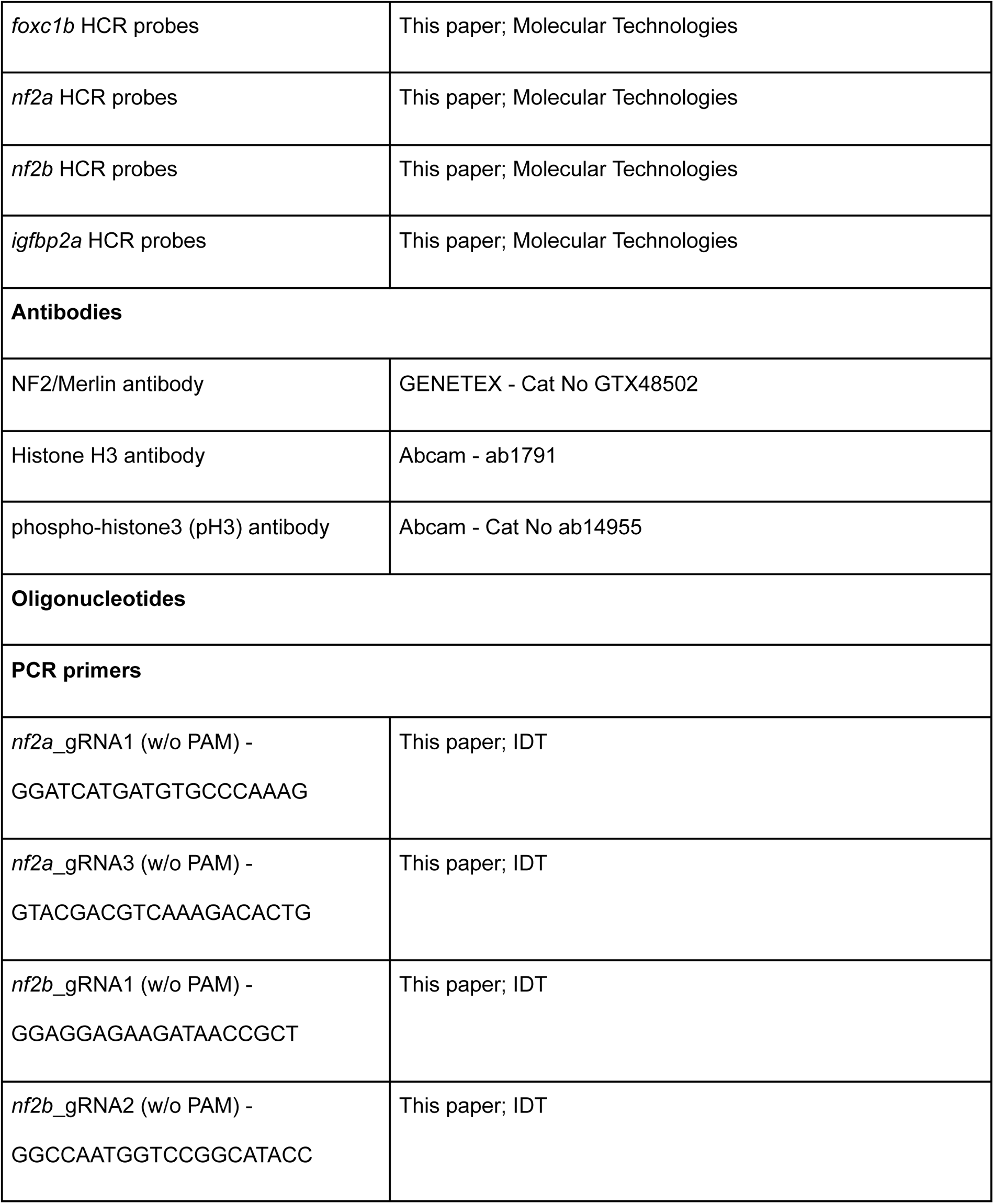
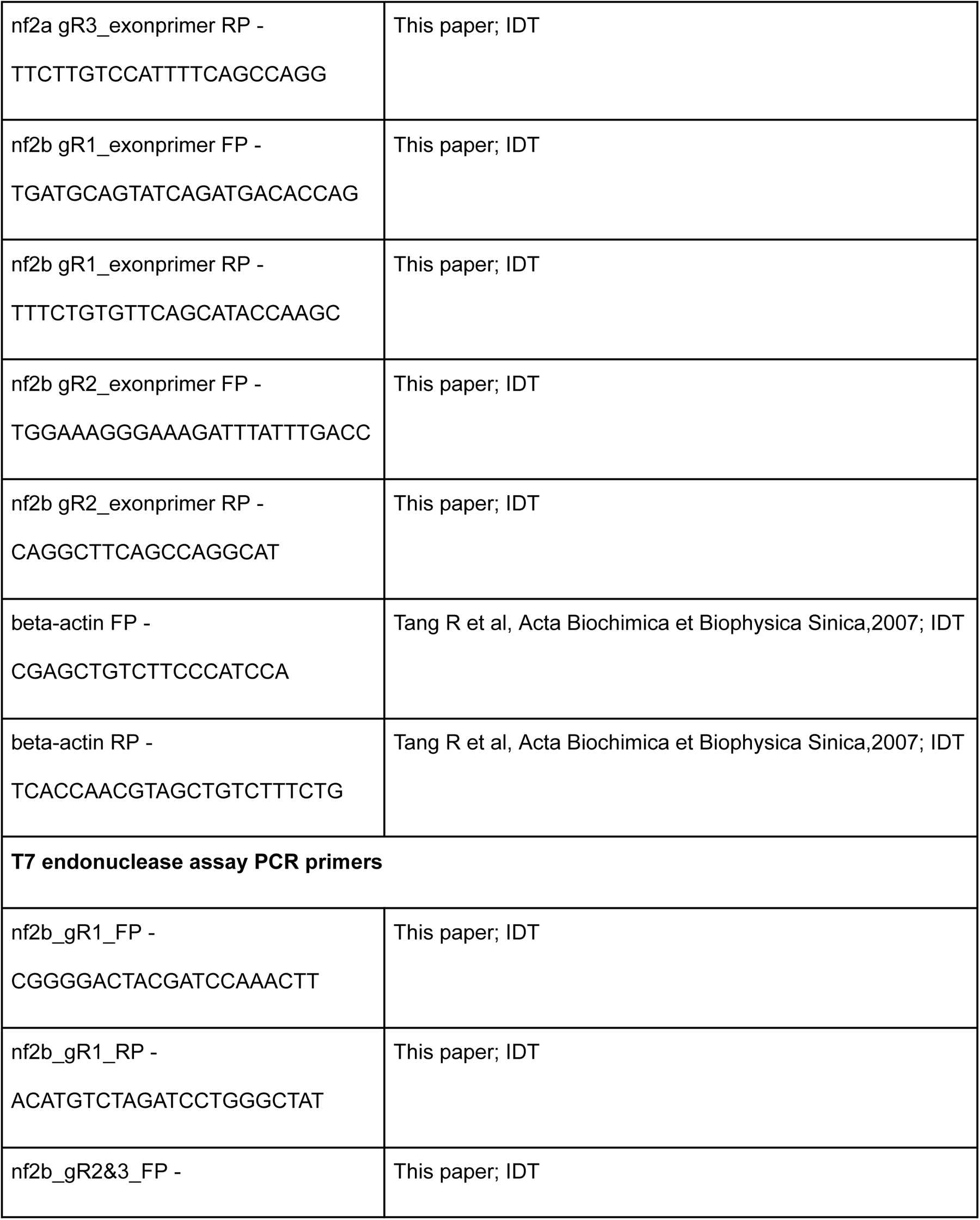
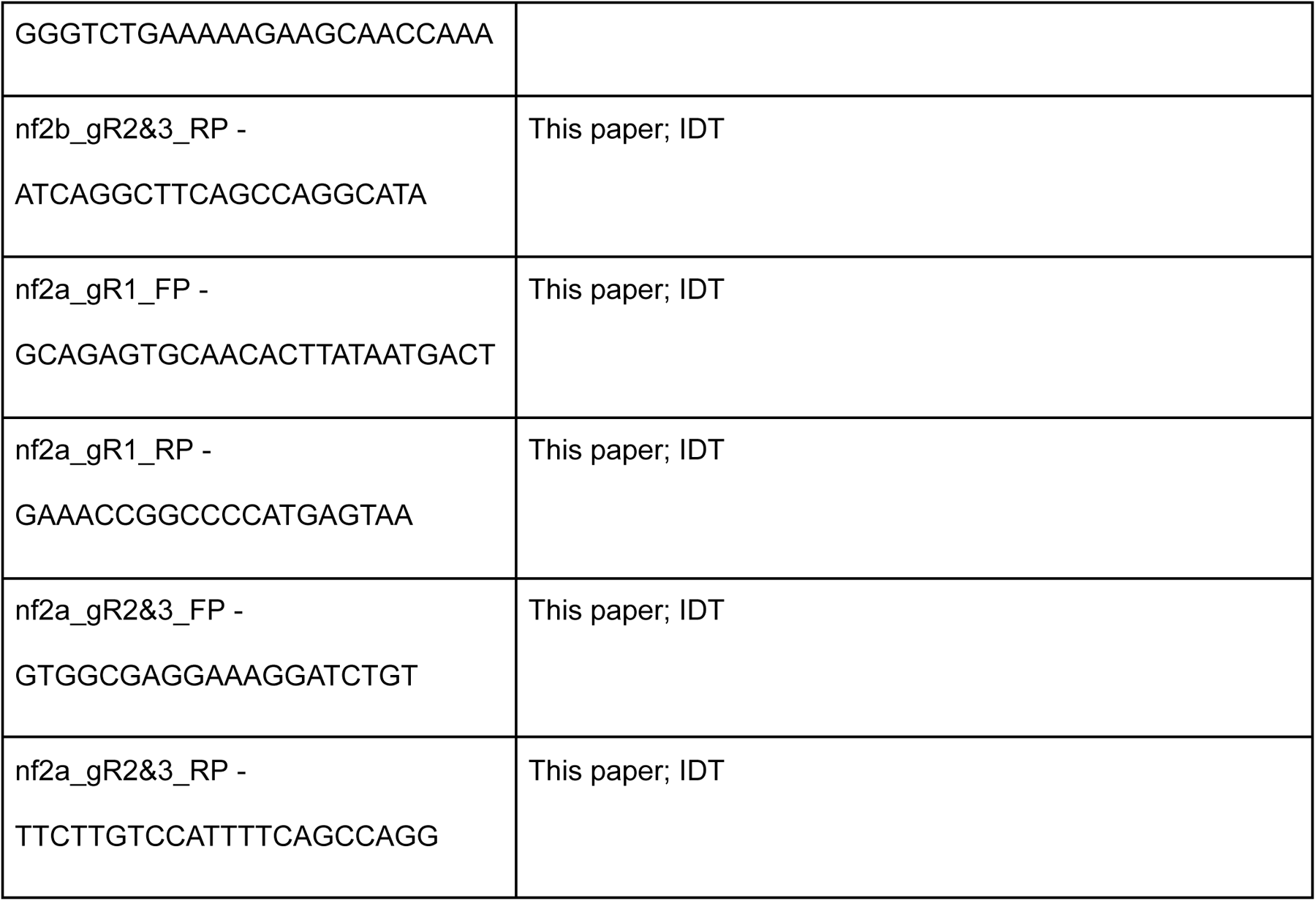

**Suppl Fig 1:**
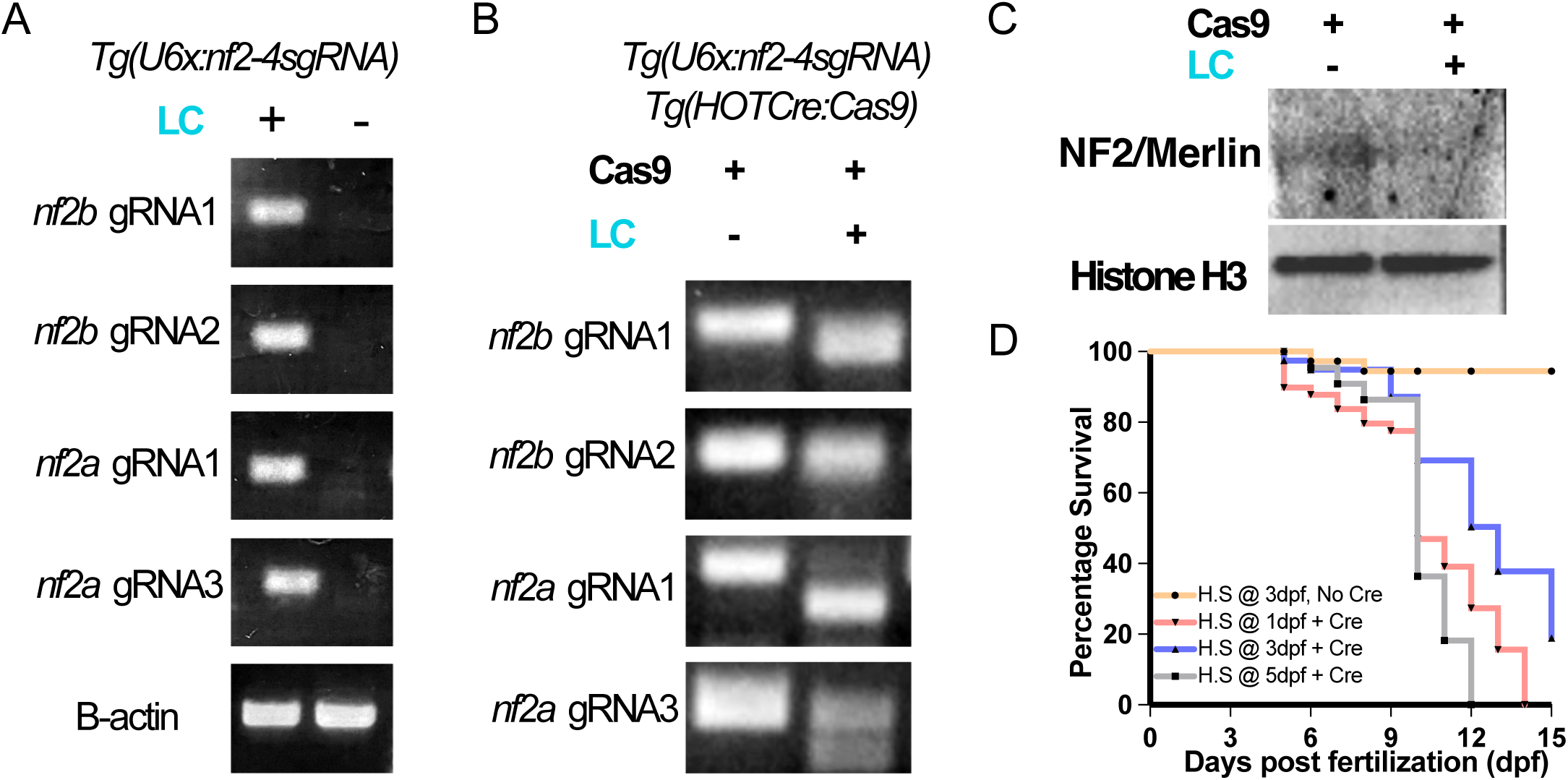
Validation of stable transgenic line *Tg(U6x:nf2-4sgRNA)::HOTCre:Cas9.* (A) Agarose gel images showing the expression of the guide RNAs in the stable transgenic line. (B) Agarose gel image of T7 endonuclease assay for *nf2* guide RNA target regions. (C) Western blot image showing down regulation of NF2 protein in *nf2* knockouts, Histone H3 was used as the loading control. (D) Survival plot of *Tg(U6x:nf2-4sgRNA):HOTCre:Cas9* after heatshock in the presence/absence of Cre-recombinase mRNA at different embryonic/larval stages. LC + indicates lens cerulean positive (contains *nf2* targeting guide RNAs) and Cas9 + indicates cardiac GFP positive (expresses Cas9 upon heatshock) embryos were used for the experiment.

**Suppl Fig 1:**
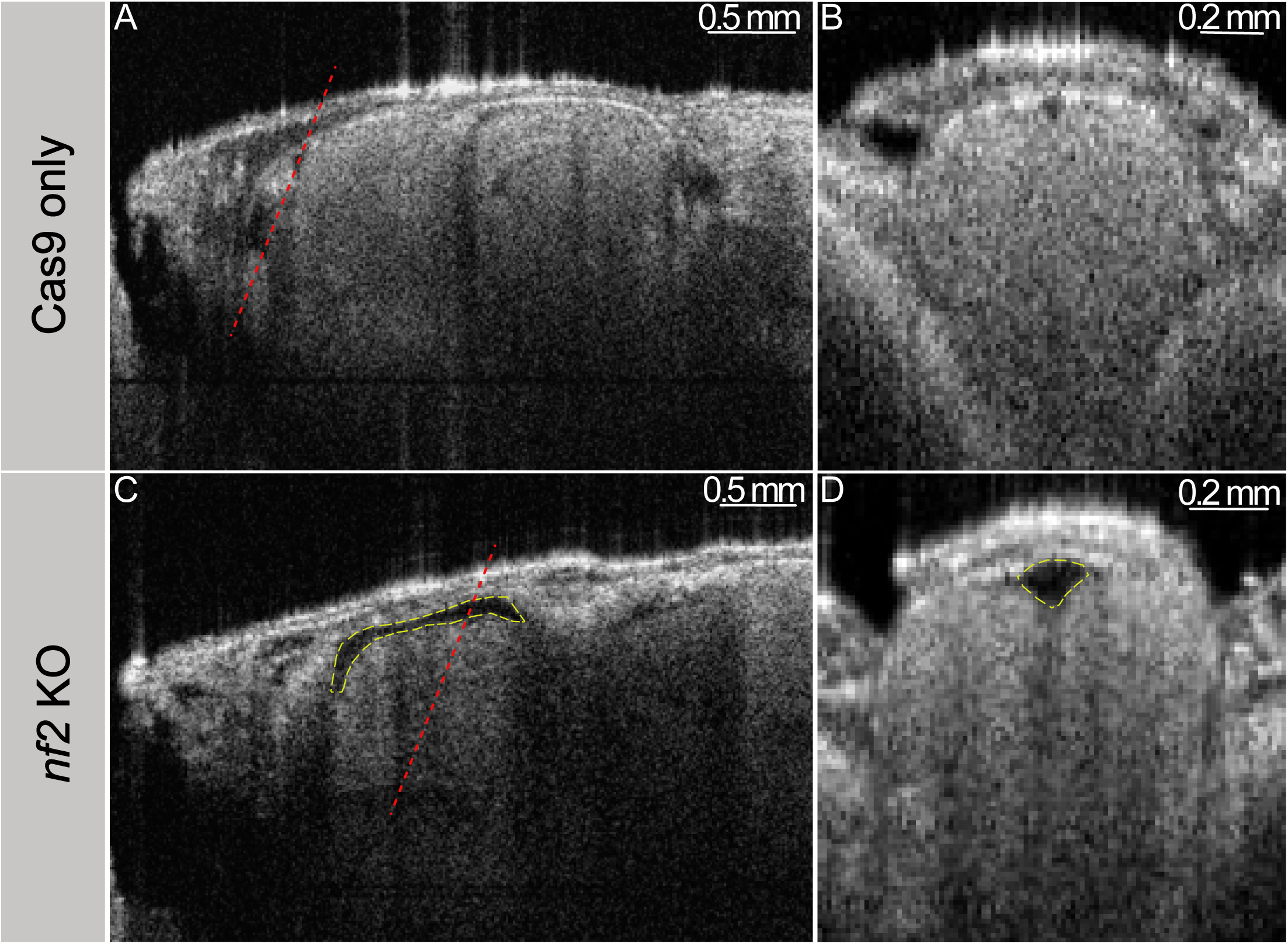
Nf2 knockout adult zebrafish display enlarged telencephalic ventricles. OCT images of euthanized adult zebrafish revealed enlarged telencephalic ventricles (highlighted with dashed yellow line) in *nf2* knockout (C,D) animals as compared to Cas9-only controls (A-B). Transverse section taken as indicated by red dashed line.

